# Host and transmission route of *Enterocytozoon hepatopenaei* (EHP) from dragonfly to shrimp

**DOI:** 10.1101/2023.02.17.528912

**Authors:** Naresh Kumar Dewangan, Jianhu Pang, Caiyuan Zhao, Changsheng Cao, Bin Yin, Shaoping Weng, Jianguo He

**Affiliations:** State Key Laboratory for Biocontrol, School of Life Sciences, Sun Yat-sen University, Guangzhou 510275, PR China, Southern Marine Science and Engineering Guangdong Laboratory (Zhuhai), Zhuhai 519000, PR China; School of Marine Sciences, Sun Yat-sen University, Zhuhai 519000, PR China

## Abstract

*Enterocytozoon hepatopenaei* (EHP) is a shrimp pathogen that causes huge economic losses. In the present study, the hosts of EHP were investigated using polymerase chain reaction (PCR) in an aquaculture farm located in Maoming, China. EHP was detected in *Litopenaeus vannamei*, *Penaeus monodon*, crab, false mussel, and three dragonfly species (*Anax parthenope*, *Pantala flavescens*, and *Ischnura senegalensis*). In the histopathological examination using hematoxylin–eosin staining, EHP spores were found in nymphs and adult dragonflies naturally infected with EHP that were collected from the shrimp farm. Fluorescence in situ hybridization results showed a positive signal for EHP infection in the fat body of dragonfly nymphs. Immature and mature microsporidian spores and late sporogonial plasmodium were observed in the cytoplasm of dragonfly nymphs using transmission electron microscopy. The transmission of EHP from shrimp to dragonfly nymphs was confirmed via cohabitation challenge experiments in which EHP-free dragonfly nymphs were cohabited with EHP-infected shrimp, and the transmission of EHP from dragonfly nymphs to shrimp was demonstrated via the cohabitation of EHP-infected dragonfly nymphs with EHP-free shrimp and oral administration challenge experiments. This study confirms that dragonflies can act as natural EHP hosts, and a novel EHP horizontal transmission route exists between dragonflies and shrimp.

**Author summary:** To the authors’ knowledge, this study presents the first report of microsporidia (EHP) infecting both crustaceans and insects (*A. parthenope*, *P. flavescens*, and *I. senegalensis*). The horizontal transmission of EHP between dragonfly nymphs and shrimp was confirmed through cohabitation and oral administration challenge experiments. EHP has become a globally significant threat to shrimp aquaculture. The findings of the present study will help to design prevention strategies, such as the use of nets to prevent dragonflies from entering shrimp ponds.

## Introduction

The microsporidian *Enterocytozoon hepatopenaei* (EHP) is an intracellular parasite that has become a serious threat to cultured shrimp in Asia, particularly in China, Vietnam, Thailand, Indonesia, India, and Malaysia ^[1, 2]^. EHP was first reported in 2004 in the hepatopancreas (HP) of growth-retarded *Penaeus monodon* in Thailand, and subsequently named and described in detail in 2009 ^[3]^. Affected shrimps suffer significant growth retardation and low-level mortality during the culture period. Often, farmers fail to notice EHP-related signs and continue to provide feed and other inputs as usual, leading to substantial economic loss (Patil et al. 2021).

EHP belongs to the genus *Enterocytozoon*, phylum Microsporidia. EHP forms single-nucleus, oval-shaped spores (1.1 μm × 0.7 μm) with five to six coils of the polar filament (PF) at one end and an anchoring disk (AD) at the other ^[3]^. Approximately 187 genera and over 1,400 species of Microsporidia have been described, among which almost half infect aquatic species and approximately 50 genera potentially infect aquatic arthropods ^[4, 5]^. Microsporidians can infect a wide variety of animals ranging from invertebrates to vertebrate hosts, including humans and arthropods like insects and crustaceans ^[6]^. Many microsporidia, including *Systenostrema alba*, *Amblyospora connecticus*, *Paranosema locustae*, *Brachiola algerae*, *Nosema bombycis*, and *Nosema carpocapsae*, have been reported in insects, but EHP has not been previously reported in insects according to the best of our knowledge ^[7–9]^. Microsporidia have different transmission strategies, including horizontal transmission (the oral transmission of spores through contaminated food and water) and vertical transmission (from parents to offspring) ^[10]^.

In recent years, whiteleg shrimp (*Litopenaeus vannamei*) have been farmed on a large scale due to their excellent traits ^[11]^. Today, *L. vannamei* account for roughly 70% of the world’s cultured shrimp. Despite the widespread adoption of specific-pathogen-free (SPF) shrimp post-larvae (PL), the fact that EHP infections are still prevalent in major shrimp-farming areas around the world ^[12]^ serves as a reminder that there may be other alternative organisms that serve as hosts for EHP transmission in the shrimp culture environment. Although there have been reports of polychaetes, *Artemia*, false mussels (*Mytilopsis leucophaeata*), and other organisms used as shrimp feed that have tested positive for EHP ^[13–16]^, it is difficult to distinguish between passive (mechanical) carriers or hosts ^[17]^. In the current critical situation of a widespread EHP epidemic, it is imperative to identify which organisms are likely to transmit EHP, especially those that inhabit the same environment as cultured shrimp.

This study investigated the host range of EHP in shrimp culture ponds with an outbreak of EHP infection and identified a new horizontal transmission mode of EHP between dragonfly nymphs and shrimp. This provides a new understanding of the host range and transmission routes of EHP and demonstrates that EHP can infect insects across species, providing new insight into its prevention and control in shrimp farming.

## Results

### Hosts of EHP

Nested PCR was used to identify the hosts of EHP in 1110 samples that included crustaceans, insects, mollusks, and fish collected from ponds with an outbreak of EHP infection in cultured shrimp. EHP was detected in *L. vannamei*, *P. monodon*, crab, three species of dragonflies, and false mussels. Out of a total of 747 samples of *L. vannamei*, 622 tested positive for EHP using nested PCR, with an infection rate of 83%, compared to 38 of 50 samples of *P. monodon* infected with EHP, with an infection rate of 73%. Four and six EHP-positive samples were detected in 13 crabs and 12 false mussels, respectively. EHP was detected in both nymphs and adults of three species of dragonfly (Table 2).

Three species of dragonfly were taken for molecular biology identification of the species and amplified with mitochondrial cytochrome oxidase subunit I (COI) primers (LCO1490 and HC02198) (S Figure 1). The sequencing results showed 99% homology to GenBank records for *A. parthenope* (MT408029) (Figure 1A&B), *I. senegalensis* (MT506706) (Figure 1C&D), and *P. flavescens* (KU641604) (Figure 1E&F).

**Figure 1.**
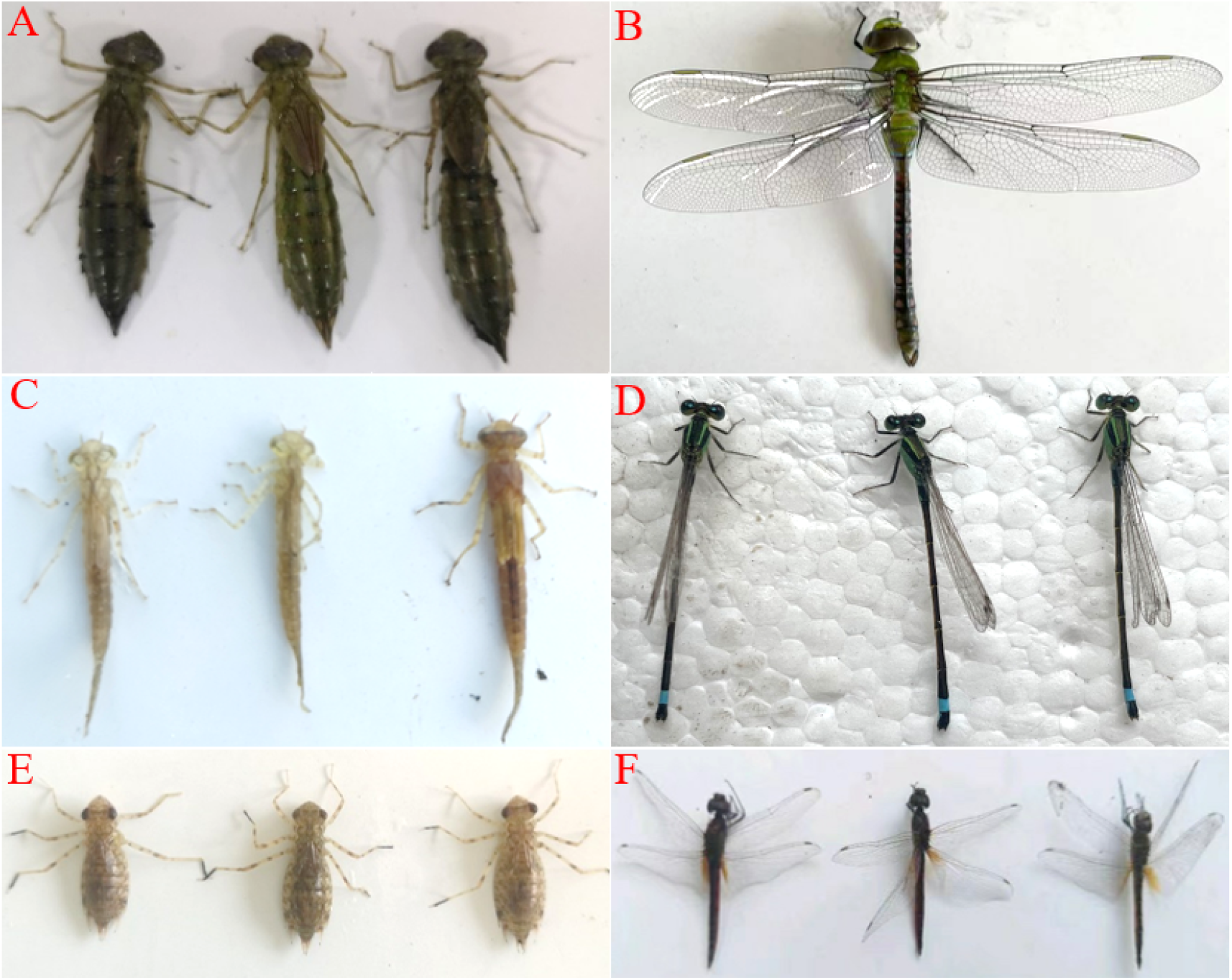
Three species of dragonflies. A: *Anax parthenope* nymphs; B: *Anax parthenope* adult; C: *Ischnura senegalensis* nymphs; D. *Ischnura senegalensis* adults; E. *Pantala flavescens* nymphs; F: *Pantala flavescens* adults.

A total of 31 *A. parthenope* nymph samples and 26 *A. parthenope* adult samples were tested for EHP using nested PCR, of which eight nymph samples and nine adult samples were positive for EHP. Forty-two *I. senegalensis* nymph samples and 30 *I. senegalensis* adult samples were tested for EHP using nested PCR, of which 10 nymph samples and 14 adult samples were positive for EHP. Fifty *P. flavescens* nymph samples and 32 *P. flavescens* adult samples were tested for EHP using nested PCR, and 15 and 12 samples were positive for EHP, respectively.

### EHP transmitted from shrimp to dragonfly nymphs by cohabitation challenge

Sixty EHP-free dragonfly nymphs were cohabitated with 15 EHP-infected shrimps. Twelve dragonfly nymphs were sampled on day 10 and day 20 for EHP detection using PCR. Among the 12 samples obtained on day 10, EHP was not detected using first-step PCR, while nine EHP-positive samples were detected using second-step PCR (Figure 2, day 10). Out of 12 samples collected on day 20, one EHP-positive sample was detected using first-step PCR, while all samples tested positive for EHP using second-step PCR (Figure 2, day 20). A BLASTN search of the sequences generated using second-step PCR revealed 99% identity with microsporidia EHP sequences. On days 10 and 20, samples were obtained from the negative control group without cohabitation challenge and tested negative for EHP using first-step and second-step PCR tests. The cohabitation challenge experiment showed that EHP was transmitted from shrimp to dragonfly nymphs.

**Figure 2.**
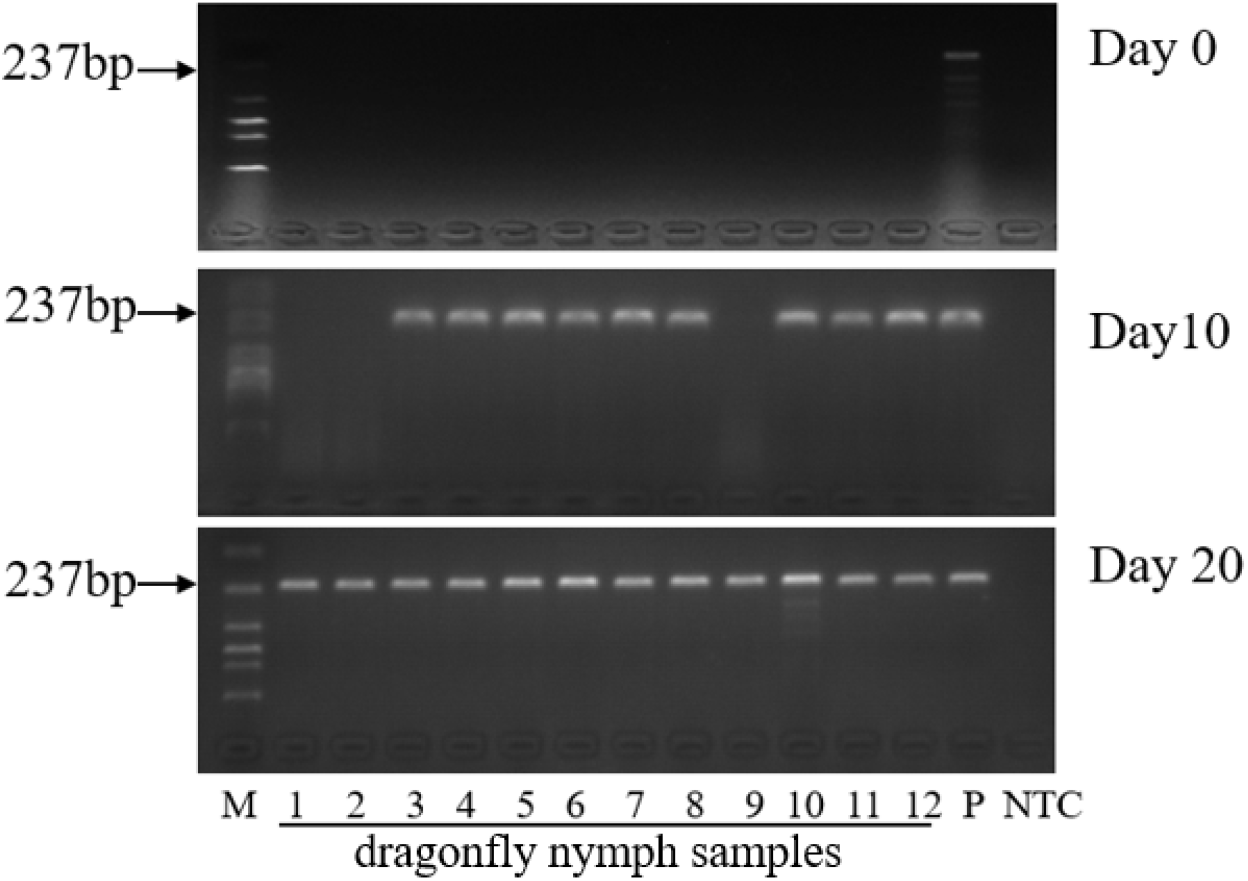
Agarose gel electrophoresis using nested PCR detection of the dragonfly nymph samples collected on days 0, 10, and 20 in the cohabitation experiment between *Enterocytozoon hepatopenaei* (EHP)-infected shrimp and EHP-free dragonfly nymphs. EHP was not detected using first-step PCR and second-step PCR in 12 dragonfly nymph samples before the start of the challenge assay (day 0); on day 10, nine out of 12 dragonfly nymph samples tested positive using second-step PCR; and on day 20, all 12 dragonfly nymph samples tested positive using second-step PCR (Lanes 1–12). Lane M: 2000-bp marker; Lane NTC: NTC using DNA extracted from dragonfly nymphs without cohabitation challenge as a template; Lanes 1–12: 12 dragonfly nymph samples collected on days 10 and 20; Lane P: positive control using DNA extracted from EHP-infected shrimp as a template. The expected amplicon size for the second-step PCR is 237 bp.

### EHP transmitted from dragonfly nymphs to shrimp by cohabitation and oral administration challenge

Thirty-six EHP-infected dragonfly nymphs from the first cohabitation challenge experiment were washed three times with sterile water and cohabitated with 40 EHP-free shrimps. This experiment was used to determine whether EHP could be transmitted from dragonfly nymphs to shrimps. Twelve shrimps were sampled from the cohabitation challenge group on days 10 and day 20 for EHP detection using PCR. Among the 12 shrimps obtained on the 10th day of the cohabitation challenge, no EHP-positive samples were detected using first-step PCR, while nine samples were found to be EHP-positive using second-step PCR (Figure 3, day 10). Twelve shrimps were obtained on day 20 of the cohabitation challenge, and four samples were found to be positive using first-step PCR, while 11 samples were found to be positive for EHP using second-step PCR (Figure 3, day 20). A BLASTN search of the sequences generated using second-step PCR revealed 99% identity with microsporidia EHP sequences. On days 10 and 20, shrimp samples were obtained from the negative control group without cohabitation challenge and tested negative for EHP using first-step and second-step PCR tests (Figure 3).

**Figure 3.**
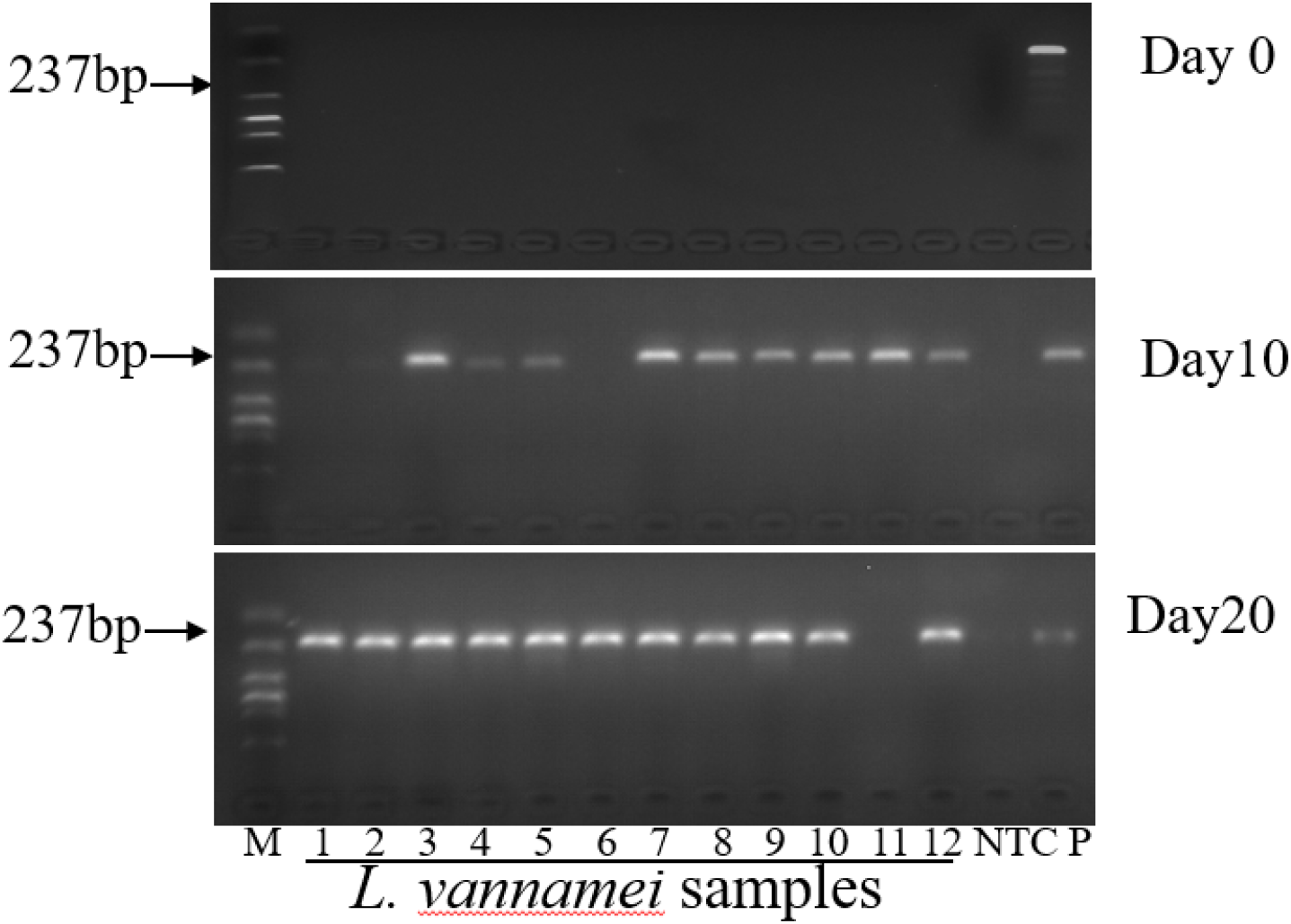
Agarose gel electrophoresis using PCR detection of the *Enterocytozoon hepatopenaei* (EHP)-free shrimp samples collected on days 0, 10, and 20 in cohabitation experiment between EHP-free shrimps and EHP-infected dragonfly nymph. EHP was not detected using first-step PCR and second-step PCR in 12 shrimp samples before the start of the challenge assay (day 0); on day 10, nine out of 12 shrimp samples tested positive using second-step PCR; and on day 20, 11 out of 12 shrimp samples tested positive using second-step PCR (Lanes 1–12). Lane M: 2000-bp marker; Lane NTC: negative control using DNA extracted from shrimps without cohabitation challenge as a template; Lane P: positive control using DNA extracted from EHP-infected shrimp as a template. The expected amplicon size for the second-step PCR is 237 bp.

Shrimps from groups 1, 2, and 4 were fed with EHP-positive *I. senegalensis* nymphs (group 1), *P. flavescens* nymphs (group 2), and EHP-infected shrimp hepatopancreas (group 4, positive control group). After 5 days of the continuous feeding of diseased tissues, three shrimp hepatopancreas samples were taken from each group on day 10 of oral administration challenge for EHP detection using PCR. Group 1, group 2, and group 4 were found to be EHP-positive using second-step PCR, while group 3, which was fed with the hepatopancreas of EHP-free shrimp, was found to be EHP-negative using the PCR test (Figure 4). The second cohabitation and oral administration challenge experiments showed that EHP was transmitted from dragonfly nymphs to shrimp.

**Figure 4.**
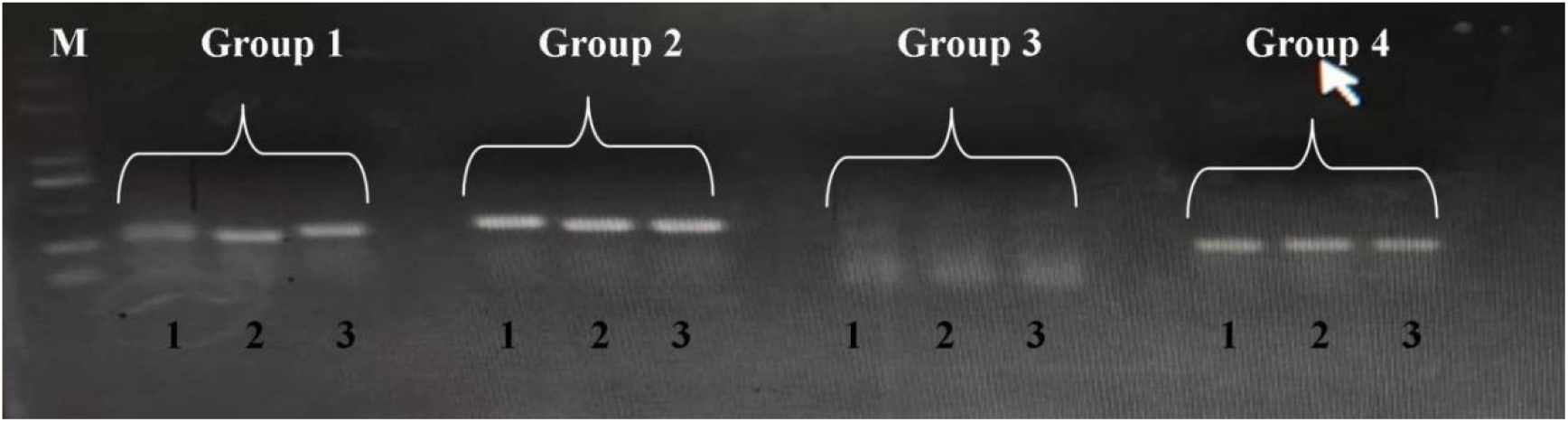
PCR result of *Enterocytozoon hepatopenaei* (EHP) in experimental shrimp. Groups 1, 2, and 4 were fed *Ischnura senegalensis* nymphs, *Pantala flavescens* nymphs, and EHP-infected hepatopancreas of shrimp, respectively, and showed EHP-positive results using second-step PCR. The size of the PCR product is 265 bp. Group, which was fed EHP-free hepatopancreas and muscle, showed negative results according to the PCR test. M=2000 bp marker.

### Histopathological examination

Histopathological examination using light microscopy revealed mature spores in the fat body of *I. senegalensis* nymphs that were cohabitated with EHP-infected shrimps in HE-stained sections (Figure 5C), while no spores were found in the intestinal and muscle tissues from the same *I. senegalensis* nymphs (Figure 5A&B). In addition, no spores were observed in the fat body tissues of EHP-free dragonfly nymphs tested using PCR (Figure 5D).

**Figure 5.**
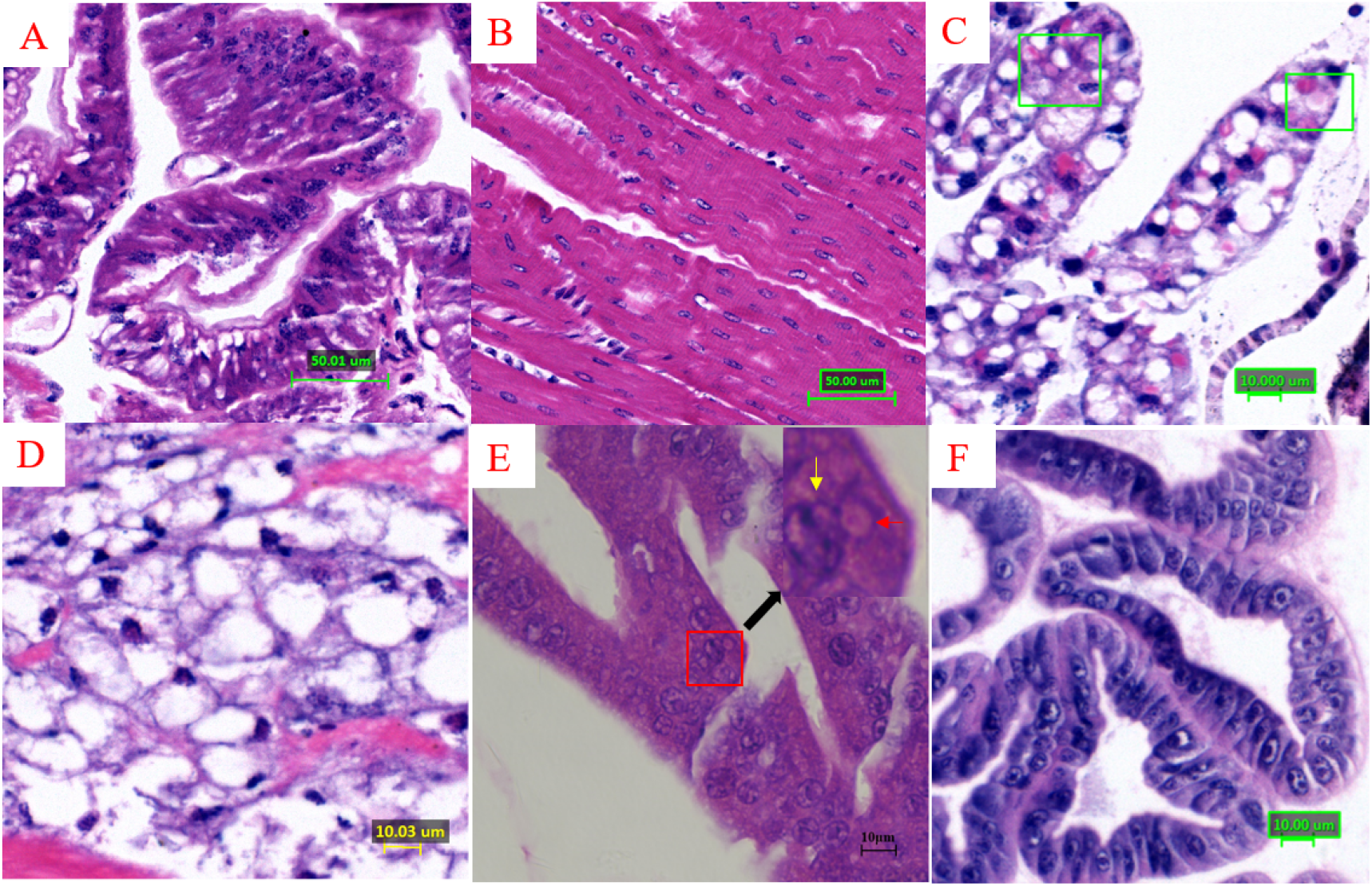
Histological examination of artificially infected *Ischnura senegalensis* nymph tissues with *Enterocytozoon hepatopenaei* (EHP). The tissues include (A): intestine, (B): muscle, (C): fat body, (D): fat body tissue of dragonfly nymphs without EHP (hematoxylin–eosin (HE) staining) and (E) & (F): hepatopancreas of shrimps. A small amount of acidophilic, granular inclusions (box) in the cytoplasm of the fat body was observed (C), and (E) acidophilic, granular inclusions (red arrow) were observed surrounding the nuclei of tubular epithelial cells (yellow arrow) of hepatopancreatic tissue from cohabitation challenge shrimps. (F): HE-stained sections of EHP-free shrimp hepatopancreas tissue.

Histopathological examination revealed acidophilic, granular inclusions in the cytoplasm of tubule epithelial cells (box) of hepatopancreatic tissue from shrimps cohabitated with EHP-infected *I. senegalensis* nymphs (Figure 5E). In contrast, histopathological examination of the hepatopancreas tissue of the EHP-free shrimp showed that the hepatopancreas tissue was intact and no microsporidian spores were observed (Figure 5F).

### FISH analysis

To determine whether specific microsporidian EHP infection was present in dragonfly tissues, a probe specific for EHP was designed. The results of FISH showed positive EHP signals (green fluorescence, white arrow) in the cytoplasm of the fat body of naturally EHP-infected and cohabitation-challenge-infected *I. senegalensis* nymphs (Figure 6C&E), while no positive signals were observed in the gut and muscle tissues from the same naturally infected dragonfly nymph samples and in the fat body tissues of EHP-free dragonfly nymphs (Figure 6 A, B&D). When the specific WSSV probe was used to detect the fat body tissues of cohabitation-challenge-infected *I. senegalensis* nymphs, no positive fluorescence signal appeared (Figure 6F). These results suggests that EHP infects dragonfly nymphs and that dragonflies are natural hosts of EHP.

**Figure 6.**
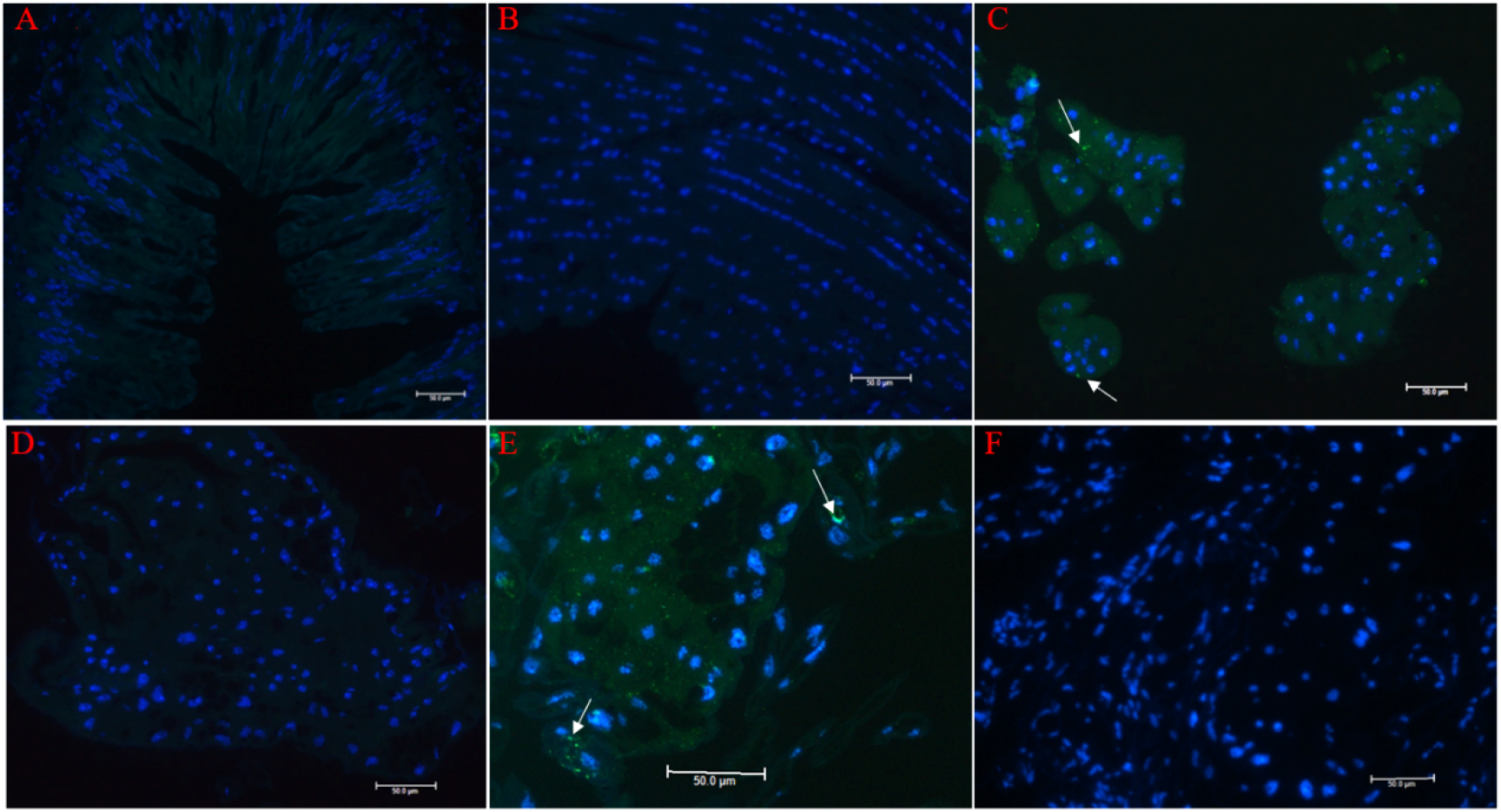
Fluorescence in situ hybridization (FISH) micrographs of various tissues of naturally infected and cohabitation-challenge-infected *Ischnura senegalensis* nymphs with *Enterocytozoon hepatopenaei* (EHP). A–C show the gut, muscle, and fat body tissues from naturally EHP-infected dragonfly nymph samples, respectively. D shows the fat body tissue from EHP-free dragonfly nymphs (negative control). E and F show the fat body tissues from cohabitation-challenge-infected *Ischnura senegalensis* nymphs with EHP. A–E: tissues of dragonfly nymphs stained with an EHP-small subunit ribosomal RNA (SSU) probe. F: tissues of dragonfly nymphs stained with a white spot syndrome virus (WSSV) probe. A, B, and D show no reaction of the EHP-SSU probe with the gut and muscle of dragonfly nymphs; only C and E (fat body) show positive signals of the EHP-SSU probe in the cytoplasm of epithelial cells of the dragonfly nymphs.

EHP-positive signals (green fluorescence, white arrow) were also observed in the epithelial cytoplasm of naturally infected shrimp hepatopancreas tissue (Figure 7 A1&B1). The FISH results of experimental challenge shrimps cohabitated with EHP-infected *I. senegalensis* nymphs (Figure 7 A2&B2) showed positive signals (green fluorescence, white arrow) for EHP in the cytoplasm of the hepatopancreatic epithelium. In contrast, the FISH results (Figure 7 A3&B3) of the negative control group, which was fed EHP-free shrimp hepatopancreas, showed no green fluorescent signal for EHP in the cytoplasm of the hepatopancreatic epithelium. These results indicate that shrimps were infected with EHP via cohabitation with dragonfly nymphs.

**Figure 7.**
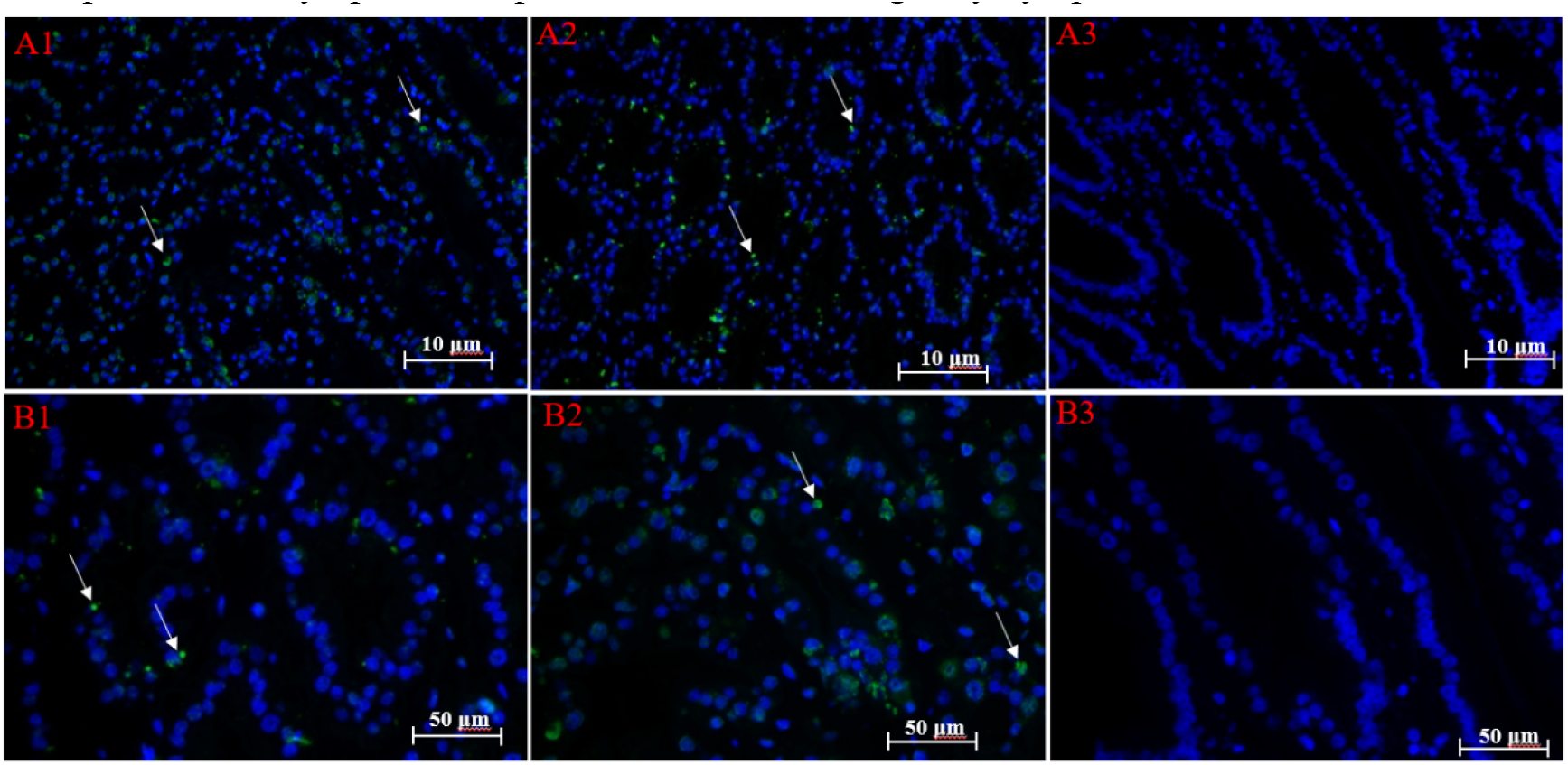
Fluorescence in situ hybridization (FISH) of hepatopancreas tissue of naturally infected and challenge-infected shrimps with *Enterocytozoon hepatopenaei* (EHP). A1&B1 show the hepatopancreas tissue of naturally infected shrimps, A2&B2 show the hepatopancreas tissue of cohabitation-challenge-infected shrimps, and A3&B3 show the hepatopancreas tissue from EHP-free shrimp samples. A1&B1 and A2&B2 show the positive signals of the EHP-small subunit ribosomal RNA (SSU) probe in the cytoplasm of epithelial cells of the hepatopancreas tissue of shrimps. A3&B3 show no positive signals for the EHP-SSU probe in the hepatopancreas epithelial cytoplasm samples of EHP-free shrimps.

### TEM observation

Immature spores (Figure 8A) and mature EHP spores (Figure 8B) were observed using TEM in the cytoplasm of cohabitation-challenge *I. senegalensis* nymphs, and four PF coils were clearly observed in mature spores (Figure 8B). The exospore and endospore (indicated by the arrow) were clearly observed and were around 1–2 μm in diameter (Figure 8B). Late sporogonial plasmodium was also observed in the cytoplasm of cohabitation-challenge dragonfly nymphs (Figure 8C).

**Figure 8.**
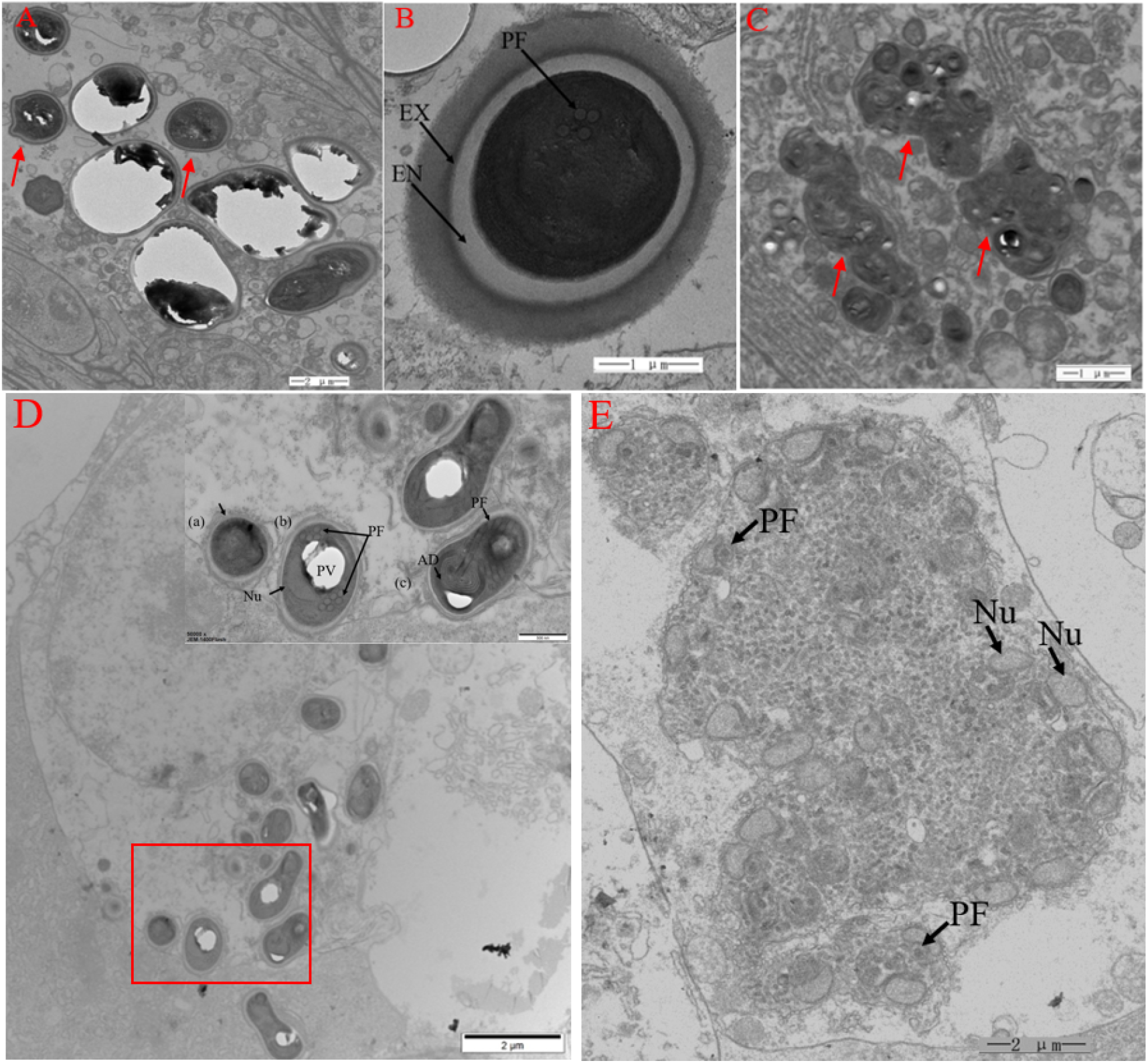
*Enterocytozoon hepatopenaei* (EHP) spores observed using transmission electron microscopy (TEM) in the cytoplasm of cohabitation-challenge *Ischnura senegalensis* nymphs and shrimps. PF: Polar filament; EX: Exospore; EN: Endospore. Immature spore (A), mature spore (B), and late sporogonial plasmodium (C) in the cytoplasm of the cohabitation-challenge *Ischnura senegalensis* nymphs. (D) Several spores can be seen in the cytoplasm of the hepatopancreatic epithelium of cohabitation-challenged shrimps. (a) Immature spores are covered with the plasmodial membrane (arrow). (b) Mature spore showing the PF, posterior vacuole (PV), nucleus (Nu), and the coiled portion of the PF. (c) Cross section of a mature spore just below the anchoring disk (AD) showing a cross section of the PF surrounded by the lamellar portion of the polaroplast and a longitudinal section of the PF outside the polaroplast. (E). Early sporogonial plasmodium containing multiple Nus and precursors of PFs in hepatopancreatic tubule epithelial cells of cohabitation-challenged shrimps.

Immature spores (Figure 8D) and mature spores (Figure 8D) were observed using TEM in the cytoplasm of hepatopancreatic epithelium from shrimps cohabitated with EHP-infected *I. senegalensis* nymphs. Immature spores were covered with the plasmodial membrane (arrow) (Figure 8D). In mature spores, the PF, the posterior vacuole (PV), the nucleus (Nu), and the coiled portion of the PF were observed (Figure 8D). A cross section of a mature spore just below the AD showed a cross section of the PF, which was surrounded by the lamellar portion of the polaroplast, with a longitudinal section of the PF outside the polaroplast (Figure 8D). Multinucleated early sporogonial plasmodium was also observed using TEM in the cytoplasm of hepatopancreatic tubules from cohabitation-challenge shrimps (Figure 8E).

## Discussion

Several microsporidian species found in arthropods mostly infect insects, and only a small number of these parasites have been detected in well-studied crustacean species ^[5]^. Microsporidia severely impact beneficial insects such as silkworms and honeybees (Becnel and Andreadis, 1999; Nageswara *et al*. 2004). This microsporidian EHP was previously reported mainly in shrimps, including *L. vanname*i and *P. monodon*. In the present work, three dragonflies (*A. parthenope*, *P. flavescens*, and *I. senegalensis*) were found to be natural hosts of EHP and not mechanical carriers of EHP, like false mussels ^[16]^. It is still unknown whether EHP can be transmitted between dragonfly and shrimp. To the best of our knowledge, there have been no reports of a microsporidia species that can infect both insects and crustaceans. Although there have been some reports of microsporidia infection in insects of the Odonata ^[9, 21]^, *A. parthenope*, *P. flavescens*, and *I. senegalensis* have not been reported to be infected with microsporidia other than the microsporidian EHP found in the present study.

*A. parthenope*, *P. flavescens*, and *I. senegalensis* are widely distributed in the tropics and subtropics. The nymphs of these species live in a variety of freshwater and brackish waters, including lakes, bogs, seepages, rivers, and springs ^[22, 23]^. In this study it was observed that many adult dragonflies laid their eggs in shrimp ponds during the period from June to October each year, which indicated that dragonfly nymphs were indeed adaptable to the environment of the shrimp culture water. The survey also found that some dragonfly nymphs collected from naturally infected EHP shrimp ponds were positive for EHP. During the cohabitation and oral administration challenge, it was observed that dragonfly nymphs fed on dead shrimp. Based on these observations and the results of the experiments, dragonflies may play a major role in the epidemiology of EHP.

Most of the previously reported microsporidia in insects can infect their fat bodies, which serve as the main organ for biosynthesis and energy storage in the insect, as well as the central tissue for metabolic activities such as growth, development, metamorphosis, and reproduction ^[24]^. The fat body function in insects is similar to that of the hepatopancreas of shrimp ^[25]^. The microsporidium *Nosema bombycis* mainly infects the fat body, midgut, silk gland, and gonad of the silkworm ^[7]^. *Paranosema locustae*, an entomopathogen of grasshoppers and locusts that mainly infects the fat body tissue, remains the only microsporidium registered and available for long-term pest control ^[8]^. An octospore microsporidium found in nymphs of *Aeshna viridis* collected from seasonal rivers ^[9]^ exhibited limited infection in the fat body and resulted in the formation of white cysts containing mature octospores. Similarly, the results of the present study showed that EHP can infect the fat body of dragonfly nymphs.

Despite widely applied control measures, including the use of EHP-free SPF shrimp larvae and the control of monodon slow growth syndrome (MSGS) by pH adjustment in shrimp culture ^[1]^, EHP is still widely prevalent in major shrimp culture regions worldwide, leading to huge economic losses. Therefore, based on the results of this research, this study proposes a new EHP prevention and control strategy to prevent dragonflies from entering the culture environment during shrimp culture and to cut off the transmission pathway of EHP.

## Materials and Methods

### Sampling

Sampling was conducted in a shrimp culture farm (Guangdong Guanlida Marine Biology Co., Ltd.) of Guangdong province of China in October and November 2019, and from July to October 2021. To investigate the host range of EHP in shrimp culture ponds with the outbreak of EHP infection, 1,110 samples including crustaceans, insects, mollusks, and fishes were collected. Crustaceans included *L. vannamei, P. monodon, Gammarus roeseli*, and crab, with 747, 50, six, and 13 samples, respectively. Insects included *Anisops kuroiwae*, *Argyroneta aquatica*, *Anax parthenope*, *Ischnura senegalensis*, *Pantala flavescens*, diving beetle, ant, and cricket, with six, six, 56, 72, 82, 11, eight, and eight samples, respectively. The samples also included *Oncomelania sail*, oyster, false mussel, and *Mugilogobius chulae* with four, 10, 12, and 19 samples, respectively. Dragonfly nymphs were collected using plankton nets, while adult dragonflies were captured from the farm area using a net. The DNA of each sample was extracted according to the method described below.

### DNA extraction and polymerase chain reaction (PCR) detection

DNA from each sample (from the hepatopancreas tissue of shrimp and the abdomen of dragonfly) was extracted using the Tissue DNA kit (Omega, Norcross, GA, USA) according to manufacturer’s instructions. The extracted DNA quantity and quality were then measured using the NanoDrop Lite spectrophotometer (Thermo Scientific, Madison, WI, USA) and stored at −20°C until use.

The detection of EHP was conducted using the nested PCR method described by Han and Mai ^[14, 18]^. The two sets of primer sequences are shown in Table 1. The PCR reaction system (25 μL) for both the first and second PCR steps contained 1x Master Mix (AG, Changsha, China) and 0.4 μM of each primer. The template for the first-step PCR contained 10 ng of DNA template, while the nested-step PCR template consisted of 1 uL amplification product from the first-step PCR. The PCR reaction procedure followed the steps published by Han and Mai. The amplicons were analyzed using 2% agarose gel electrophoresis with ethidium bromide staining and using a 2000-bp DNA ladder marker (Takara, Tokyo, Japan).

**Table 1.**
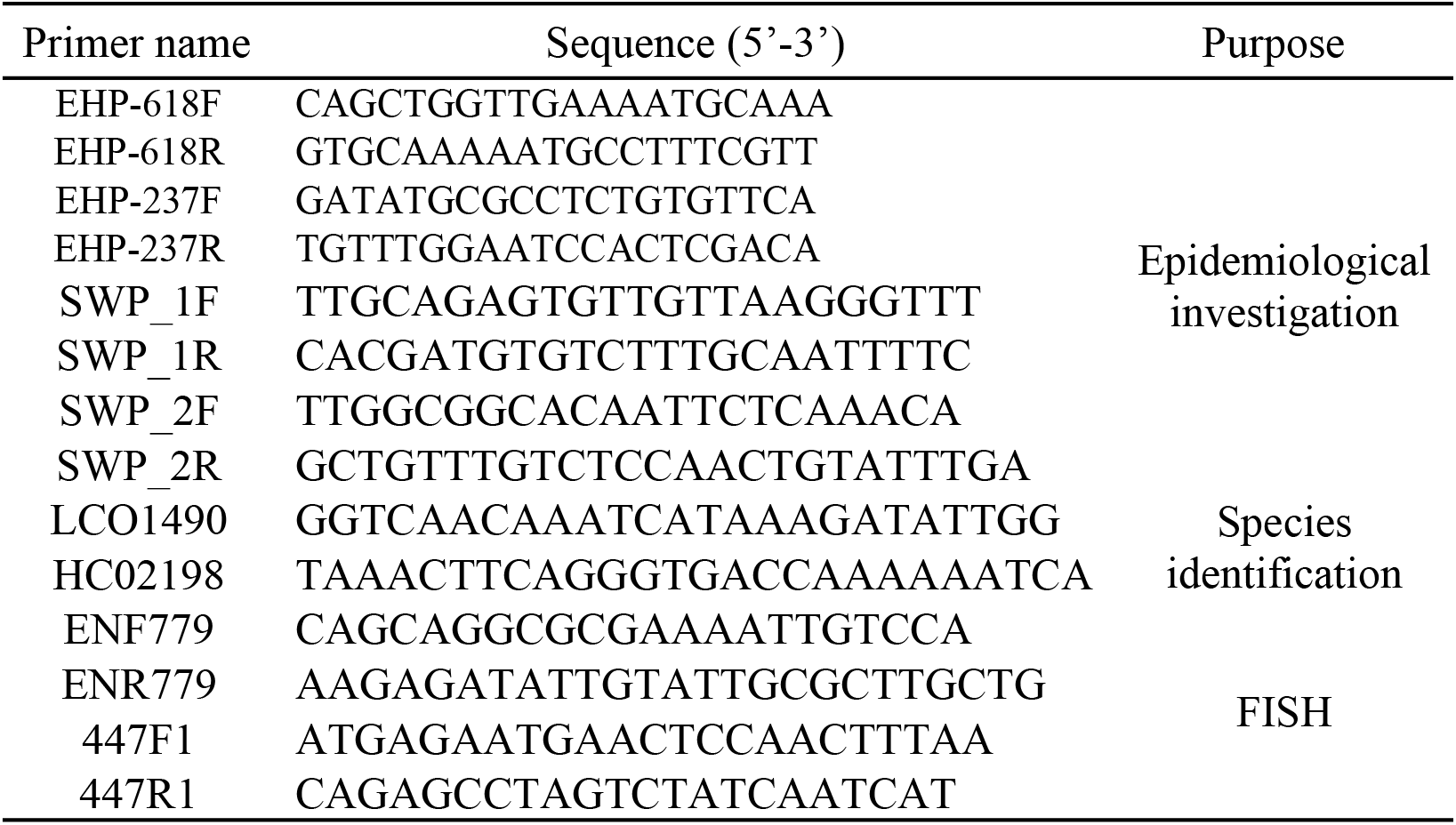
Primers used in this study

**Table 2.**
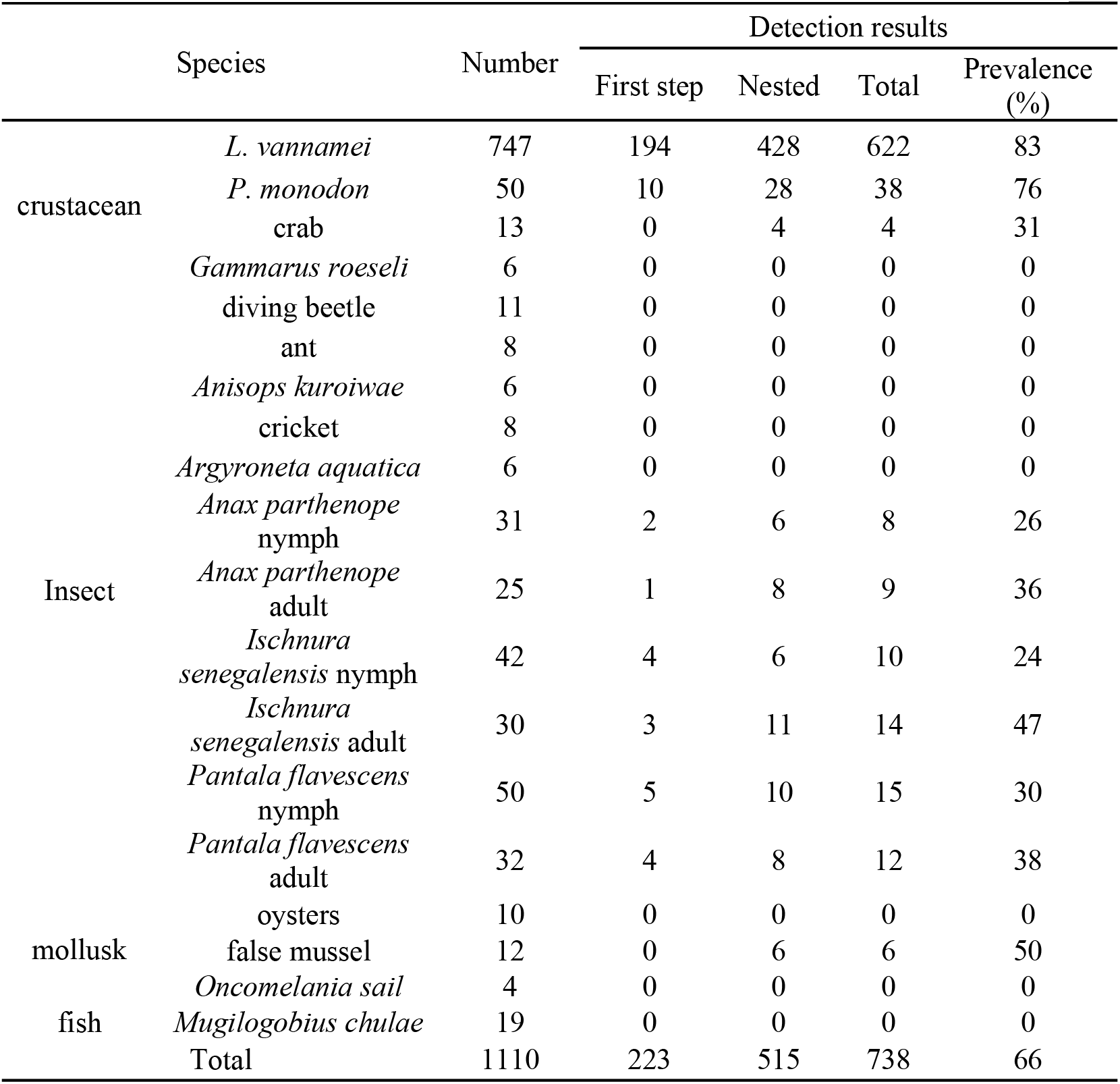
Summary of the carriers of EHP in aquatic organisms

### Molecular identification of dragonfly

The identification of dragonfly species was accomplished via molecular biology. The DNA templates of three dragonfly species (nymphs and adults) were extracted according to the above method. The mitochondrial cytochrome oxidase subunit I (COI) gene primers (LCO1490 and HC02198) (Table 1) were used for the identification of dragonfly species ^[19, 20]^. The expected amplicons of the PCR product were 710 bp, which were analyzed using 1.5% agarose gel electrophoresis with ethidium bromide staining and sequenced by Sangon (Shanghai, China).

### Cohabitation challenge

#### Cohabitation challenge between EHP-infected shrimp and EHP-free dragonfly nymphs

Prior to the start of the challenge experiment, 200 dragonfly (*I. senegalensis*) nymphs were collected from shrimp ponds without an outbreak of EHP infection. Twelve were randomly selected for EHP detection using PCR, and the first-step PCR and second-step PCR tests verified that these dragonfly nymphs were EHP-free. To confirm whether EHP could be transmitted from shrimp to dragonfly nymph, in the first cohabitation challenge experiment, 15 EHP-infected *L. vannamei* were cohabitated with 60 EHP-free dragonfly nymphs in an aquarium (50 L) that contained sufficient oxygen (dissolved oxygen >5 ppm) at a salinity of 5 ppt and a temperature of 28–32°C for 20 days. One-third of the water was changed daily and shrimp were fed twice a day, with 2% of the body weight of shrimp supplied during each feeding. Twelve dragonfly nymphs were taken from the cohabitation challenge group on days 10 and 20 for EHP detection via PCR. The amplified products were analyzed using 1.5% agarose gel electrophoresis with ethidium bromide staining and sequenced by Sangon (Shanghai, China). The remaining 36 dragonfly nymphs from this cohabitation challenge group were used for the second cohabitation challenge experiment of infecting EHP-free shrimp with the disease material. Ten dragonfly nymphs were reared under the same conditions for 20 days without a cohabitation challenge treatment as a negative control, and three samples were taken on days 10 and 20 for pool testing.

#### Cohabitation challenge between EHP-infected dragonfly nymphs and EHP-free shrimp

To confirm whether EHP could be transmitted from dragonfly nymphs to shrimp, the second cohabitation challenge experiment was performed. One hundred *L. vannamei* were collected from shrimp culture ponds and 12 were randomly selected for EHP detection using PCR. The first-step PCR and second-step PCR tests confirmed that these *L. vannamei* were EHP-free. The 36 artificially infected dragonfly nymphs that remained following the first cohabitation challenge experiment were cohabitated with 40 EHP-free *L. vannamei* under the same conditions for 20 days. The shrimp feeding amount, water change frequency, and sampling time and number were consistent with the first cohabitation experiment. Ten EHP-free shrimps were temporarily reared under the same conditions for 20 days without a cohabitation challenge treatment as a negative control, and three samples were taken on day 10 and 20 for EHP pool testing via PCR.

#### Oral administration challenge

Experimental transmission by oral administration challenge was performed in four groups, including three experimental groups and one control group. Each group consisted of two aquariums and each aquarium (50 L) contained 10 *L. vannamei* (20 shrimps in each group) reared under the same conditions for 10 days. All the groups of shrimps were starved for 24 hours before oral administration. Group 1 and group 2 shrimps were fed with chopped *I. senegalensis* nymphs and *P. flavescens* nymphs, respectively. Group 3 shrimps were fed the hepatopancreas tissues of EHP-free shrimps as the negative control group, while control group 4 was fed the hepatopancreas tissues of EHP-infected shrimp as the positive control group. After 5 days of the continuous feeding of diseased tissues, three shrimp hepatopancreas tissues were taken from each group on the 10th day of the challenge for EHP detection using PCR.

#### Histopathological examination

EHP-free and natural and artificial EHP infections of dragonfly nymph and shrimp samples were fixed in 4% paraformaldehyde/ (diethyl pyrocarbonate, DEPC) water (4% PFA/DEPC) at 4°C. After 24 hours the dissected tissues were transferred into gradient alcohol, and paraffin embedding was performed the following day. Tissues were sectioned at 3 μm using a microtome (Leica, Germany) and the sections were stained with hematoxylin–eosin (HE) and fluorescence in situ hybridization (FISH), then examined according to the method described by Lightner ^[2]^.

#### FISH analysis

The specific probe used for FISH analysis was prepared using primers ENF779 and ENR779 (Table 1) targeting the EHP small subunit ribosomal RNA (SSU rRNA) gene, which contained 6-carboxy-fluorescein (6-FAM) to substitute for a portion of the dTTP in the dNTP mix. The other specific probe for white spot syndrome virus (WSSV) was developed as a negative control DNA probe using the same 6-FAM-labelling kit with the primers 447F and 447R ^[16]^. The sizes of the SSU-EHP and WSSV probes were 779 bp and 447 bp, respectively. The FISH protocols were based on those described previously ^[20]^.

The visceral tissue of dragonfly nymphs and hepatopancreas of shrimp were fixed in 44% PFA/DEPC water for 12 hours at 4°C and then dehydrated using gradient alcohol. Paraffin embedding was performed the following day. Tissues were sectioned to 3-μm thickness and heated in an oven at 62°C for 2 hours. Sections were washed sequentially with Dewaxing Transparent Liquid, gradient alcohol, and DEPC dilution. Sections were permeabilized with 20 μg/mL proteinase K at 37°C and then washed with phosphate-buffered saline (pH 7.4). Hybridization was conducted at 42°C for 16 hours with the EHP DNA probes. Then, sections were washed with 2 × SSC, 1 × SSC, and 0.5 × SSC to remove the hybridization solution. Sections were incubated with 4′,6-diamidino-2-phenylindole (DAPI) (1:2000, Solarbio, Beijing, China) for 8 min in the dark. Finally, sections were mounted on slides and scanned with an Olympus slide scanner (Olympus, Tokyo, Japan).

#### Transmission electron microscopy (TEM) observation

Tissues (2 mm^3^) from EHP-positive *I. senegalensis* nymph specimens (n=3) and small pieces (2 mm^3^) of hepatopancreas from EHP-positive shrimp (n=3) were fixed in 2.5% glutaraldehyde in Millonig buffer followed by washing in 0.1 M sodium cacodylate buffer three times. The tissues were then stained in 0.5% aqueous uranyl acetate for 1 hour and were proceed for TEM analysis following the procedure of Tourtip et al. ^[3]^. The ultrastructural details of the spores within the plasmodium stages in the cytoplasm were observed using JEM-1400 Flash TEM (JEOL, Tokyo, Japan).

## Acknowledgments

This work was funded by the National Key Research and Development Program of China (2018YFD0900501) and the Marine Life Processes and Utilization of Biological Resources (42000-02920001).

## Author contributions

NK, JP and JH conceived this study, and SW, CZ, CC and, YB participated in its design. JP and NK carried out the experiments, and CZ, CC participated in data analysis. JP drafted the manuscript, SW and JH revised the manuscript. All authors read and approved the final manuscript before submission. All authors contributed to the article and approved the submitted version.

## Declaration of interests

The authors declare that the research was conducted in the absence of any commercial or financial relationships that could be construed as a potential conflict of interest.

